# Absence of CDK12 in oocyte leads to female infertility

**DOI:** 10.1101/2024.11.11.622910

**Authors:** Denisa Jansova, Veronika Sedmikova, Fatima J. Berro, Daria Aleskhina, Michal Dvoran, Michal Kubelka, Jitka Rezacova, Jana Rutarova, Jiri Kohutek, Andrej Susor

## Abstract

Transcriptional activity and gene expression are essential for the development of a mature, meiotically competent oocyte. We have found that the absence of cyclin-dependent kinase 12 (CDK12) in oocytes leads to complete female sterility, as there are no fully developed oocytes able to accomplish meiosis I in the ovaries. Mechanistically, CDK12 in growing oocytes controls POLII activity and maintenance of the physiological maternal transcriptome, which negatively affects protein synthesis that promotes further oocyte growth. In addition, disruption of oocyte development disrupts folliculogenesis, resulting in a premature failure phenotype without terminal folliculogenesis and ovulation. In summary, we have characterized a single master regulator of the oocyte transcriptional program and gene expression that is essential for oocyte growth and female fertility.

**Graphical Abstract:** 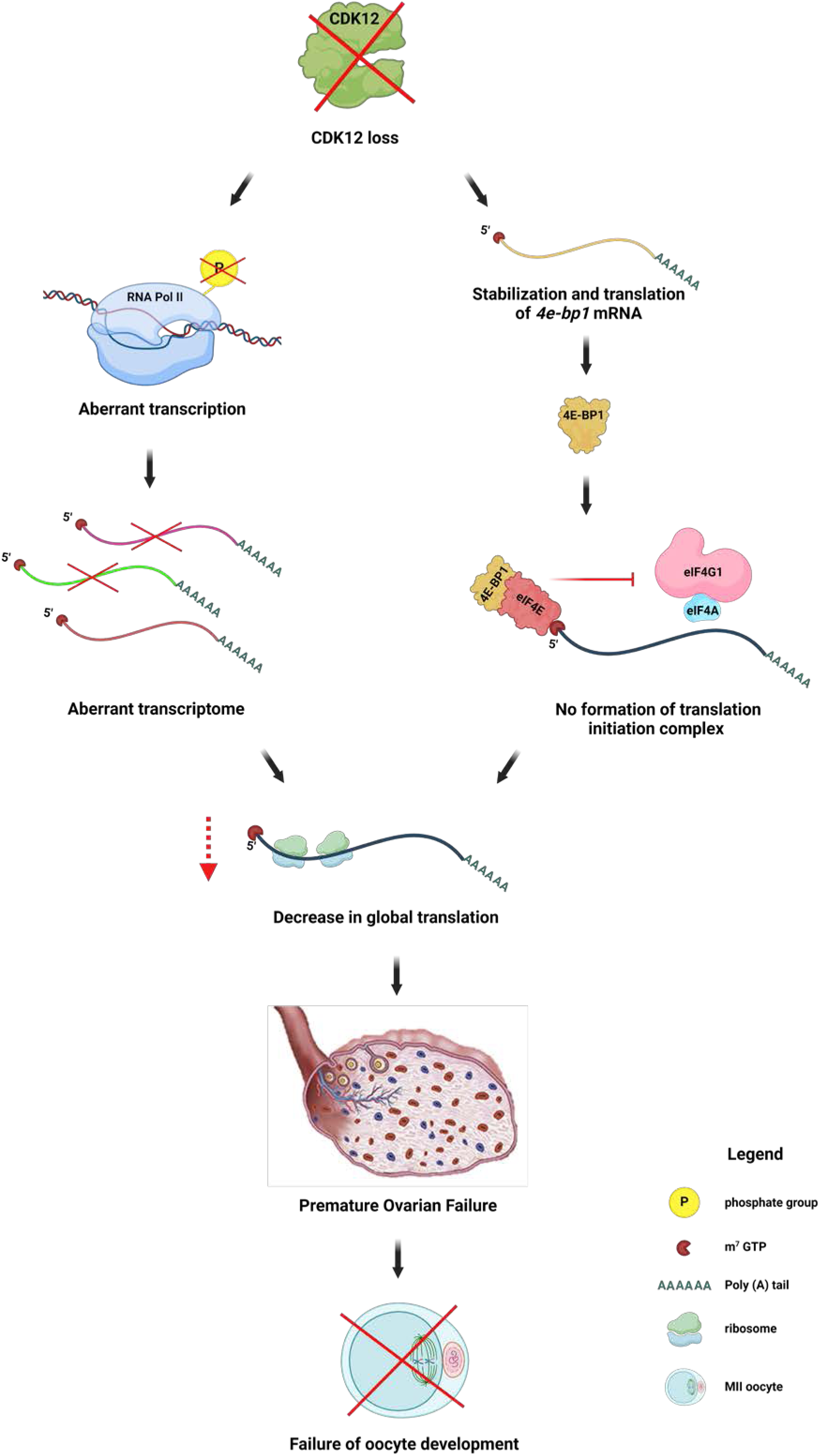

## Introduction

Oocytes arise from primordial germ cells and develop in follicles. The periodic activation of primordial germ cells up to the preovulatory stage is accompanied by oocyte growth and cytoplasmic expansion, with the synthesis and storage of maternal components such as RNA and proteins in the cytoplasm [1]. The quality of oocytes and early embryos is based on the storage of maternal components, which is regulated by various signalling pathways. During this period, the oocyte acquires meiotic competence, i.e. the ability to resume meiosis and to enter or arrest at metaphase II [2], and developmental competence, the ability to maintain early embryonic development [3].

The regulation of transcription initiation by RNA polymerase II (POLII) is of central importance for the maintenance of the oocyte and early embryonic development. Transcriptional silencing occurs in a fully-grown oocyte and continues during meiotic maturation and after fertilization in the early embryo. The hyperphosphorylated form of POLII has been found in growing oocytes, while the hypophosphorylated form is typical of mature, transcriptionally inactive GV oocytes [4]. The transcription of protein-coding genes by POLII is tightly regulated to generate proper quantities and classes of mRNA. Cyclin-dependent kinase 12 (CDK12) forms a complex with Cyclin K which controls transcription, by phosphorylating the C-terminal domain (CTD) of the large subunit of POLII [5].

Current evidence suggests that CDK12 has a wide range of biological functions, including DNA replication [6], regulation of the expression of DNA damage response genes and cell cycle genes [7], transcription elongation [8], pre-mRNA processing [9], RNA turnover [10] and the initiation of translation of mRNA subgroups [7,11,12]. Blazek *et al.* have shown that the depletion of CDK12 leads to a reduced expression of 4% of genes [5]. CDK12 is one of the most frequently mutated genes in ovarian carcinoma [13] and these mutations lead to a loss of function [14]. Although the role of CDK12 in cancer has been investigated, its role in oocyte development is unknown.

We have established an oocyte-specific knockout mouse line without expression of the CDK12 protein. In this study, we demonstrated that the absence of CDK12 impairs transcription via the regulation of POLII, resulting in a dysregulated maternal transcriptome and translation. This disruption has a negative effect on oocyte growth and thus on folliculogenesis, which ultimately leads to female infertility.

## Results

### CDK12 is essential for female fertility

To explore the requirement of Cyclin-dependent kinase 12 (CDK12) in female fertility, we performed a series of experiments using a conditional CDK12 knockout (cKO) mouse model. The aim of our study was to understand the effects of CDK12 depletion on oocyte development and overall fertility in female mice. The CDK12 cKO was generated by crossing *Cdk12^fx/fx^* with a Z*p3^Cre^* strain (**Supporting Information 1 A, B**). The resulting experimental genotypes were labelled as wild-type (WT; CDK12^+/+^; Cdk12^tm1c+/+ Zp3-Cre+/+^); heterozygote (HT; CDK12^+/-^; Cdk12^tm1c+/-Zp3-Cre^ ^+/-^) and homozygote (cKO; CDK12^-/-^; Cdk12^tm1c-/-Zp3-Cre+/+^) (**Supporting Information 1 A, B**). Immunoblot analysis confirmed the absence of CDK12 protein in the CDK12^-/-^ GV oocytes (**Fig. 1A, B** and **Supporting Information 1C, D**). In addition, immunofluorescence experiments showed that CDK12 was localized to the nucleus of WT oocytes, while it was absent in CDK12^-/-^ oocytes (**Supporting Information 1C, D**). Breeding experiments with CDK12^+/+^ proven breeder males showed that females lacking CDK12 in their oocytes were completely sterile, while CDK12^+/+^ and CDK12^+/-^ females were normally fertile (**Figure 1C**), suggesting that there is no haploinsufficiency for CDK12.

**Figure 1:**
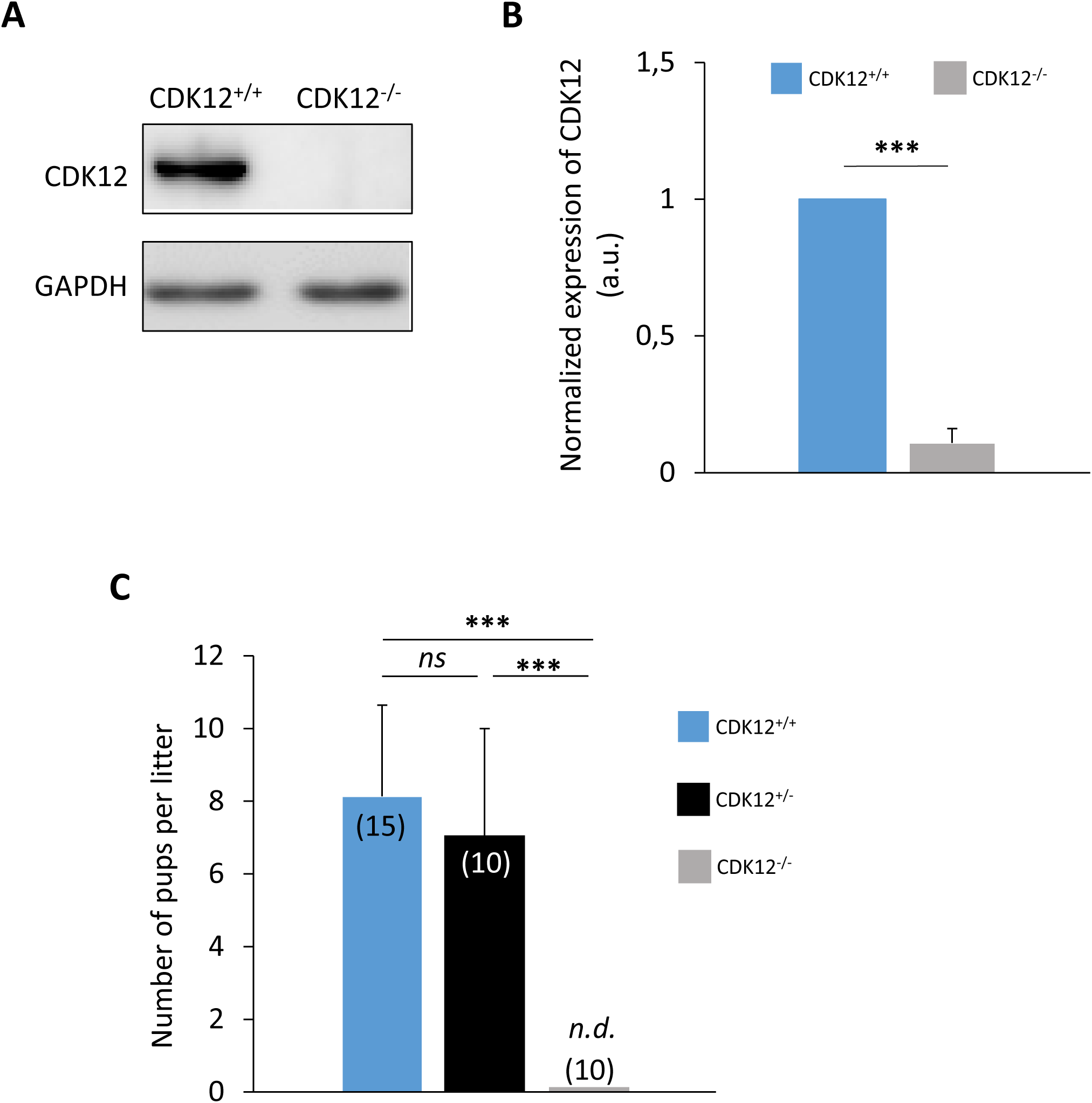
CDK12 is essential for female fertility. **A)** Western blot analysis of CDK12 expression in oocytes from wild-type (CDK12^+/+^) and homozygous conditional knockout (cKO) females (CDK12^-/-^). GAPDH was used as a loading control. Data from six independent biological replicates. For *Cdk12* mRNA expression analysis, see Fig. 5. For information about conditional KO generation and CDK12 localization and expression, see **SI** Fig. 1. **B)** Quantification of CDK12 protein expression from (**A**) normalized to GAPDH. Data are presented as mean ± SD; Student’s *t*-test: ***p < 0.001. **C)** Analysis of fertility of females from different genotypes mated with proven breeder wild type males. Number of breeding pairs in parentheses. Data are presented as mean ± SE; Student’s *t*-test: *ns*, non-significant; ***p < 0.001.

In conclusion, the absence of maternal CDK12 in oocytes leads to female sterility.

### CDK12 is Essential to Oocyte Development and Maturation

Next, we investigated why absence of CDK12 in oocytes leads to female infertility. It is known that oocytes undergo a series of developmental stages prior to fertilization, including oocyte growth, acquisition of meiotic competence and maturation into fertilizable MII oocyte. To understand the main cause of infertility in cKO mice, we analyzed the ovaries and oocyte quality. Following the standard procedure of superovulation, we examined the morphology of the ovaries. Despite having the equal body size, the ovaries of cKO females were significantly smaller compared to WT females (**Figure 2A** and **Supporting Information 2A**). Histological analysis of cKO ovaries and quantification of follicles revealed a reduced number of primary follicles and almost no antral follicles (**Figure 2B, C**), resulting in the premature ovarian failure (POF) phenotype. In addition, rare ovulations lead to the formation of the corpus luteum in the cKO ovaries, which shows that the ovaries respond to hormonal stimulation and ovulation (**Supporting Information 2B**). Next, we found that post PMSG stimulation most of the isolated oocytes from WT females were fully grown germinal vesicle (GV) (**Figure 3A, B**). In contrast, oocytes from cKO females were predominantly growing GV oocytes (**Figure 3A, B**). Following the standard procedure of superovulation, we obtained only a few ovulated oocytes in the oviduct of the CKO females, although the WT genotype ovulated a normal number of oocytes (**Figure 3B**). In contrast, cKO females do not produce MII oocytes during scarce ovulation (**Figure** 2 **3 C, D**). Instead, the ovulated CDK12^-/-^ oocytes were devoid of polar body, with disorganized chromosomes and polymerized tubulin (**Figure 3 C, D**).

**Figure 2:**
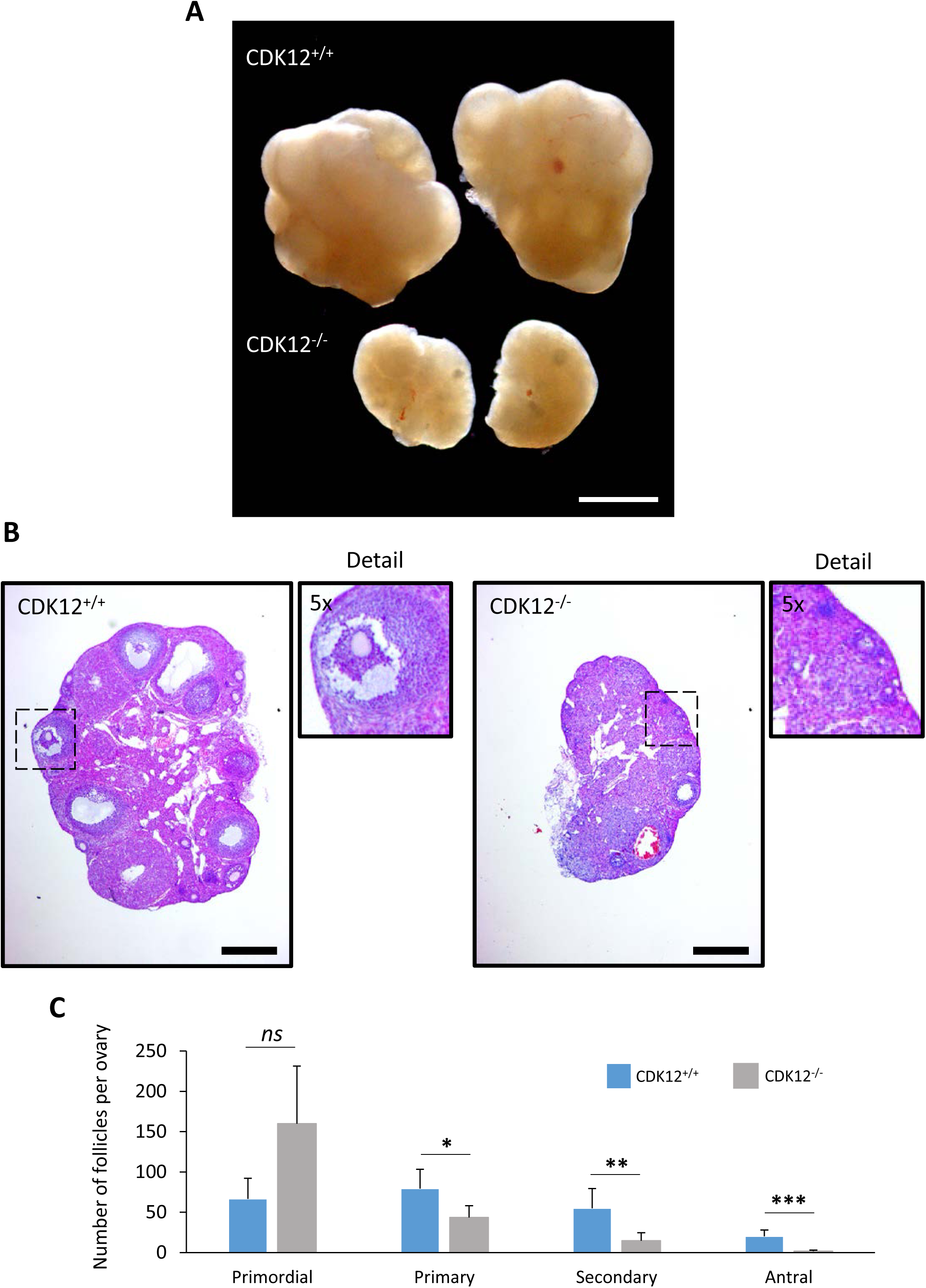
Absence of CDK12 in the oocytes leads to decreased ovarian size via ceased folliculogenesis. **A)** Representative image of pair of ovaries from CDK12^+/+^ and CDK12^-/-^ mice. Scale bar 1 mm. For measurement of the size of the ovaries, see **SI** Fig. 2. **B)** Representative images of histological sections of ovaries stained with haematoxylin and eosin. The dashed squares correspond to 5-fold magnification of the ovarian cortex, showing representative antral follicle (CDK12^+/+^) and primordial follicle (CDK12^-/-^). Data from two independent biological replicates, n= 5 per group; scale bars 400 µm. For depiction of the corpus luteum in the ovaries, see **SI** Fig. 2. **C)** Quantification of follicles from CDK12^+/+^ and CDK12^-/-^ mice. Data from three females for each genotype, data are presented as mean ± SE; Student’s *t*-test: *ns*, non-significant; *p < 0.05; **p<0.01; ***p < 0.001.

**Figure 3:**
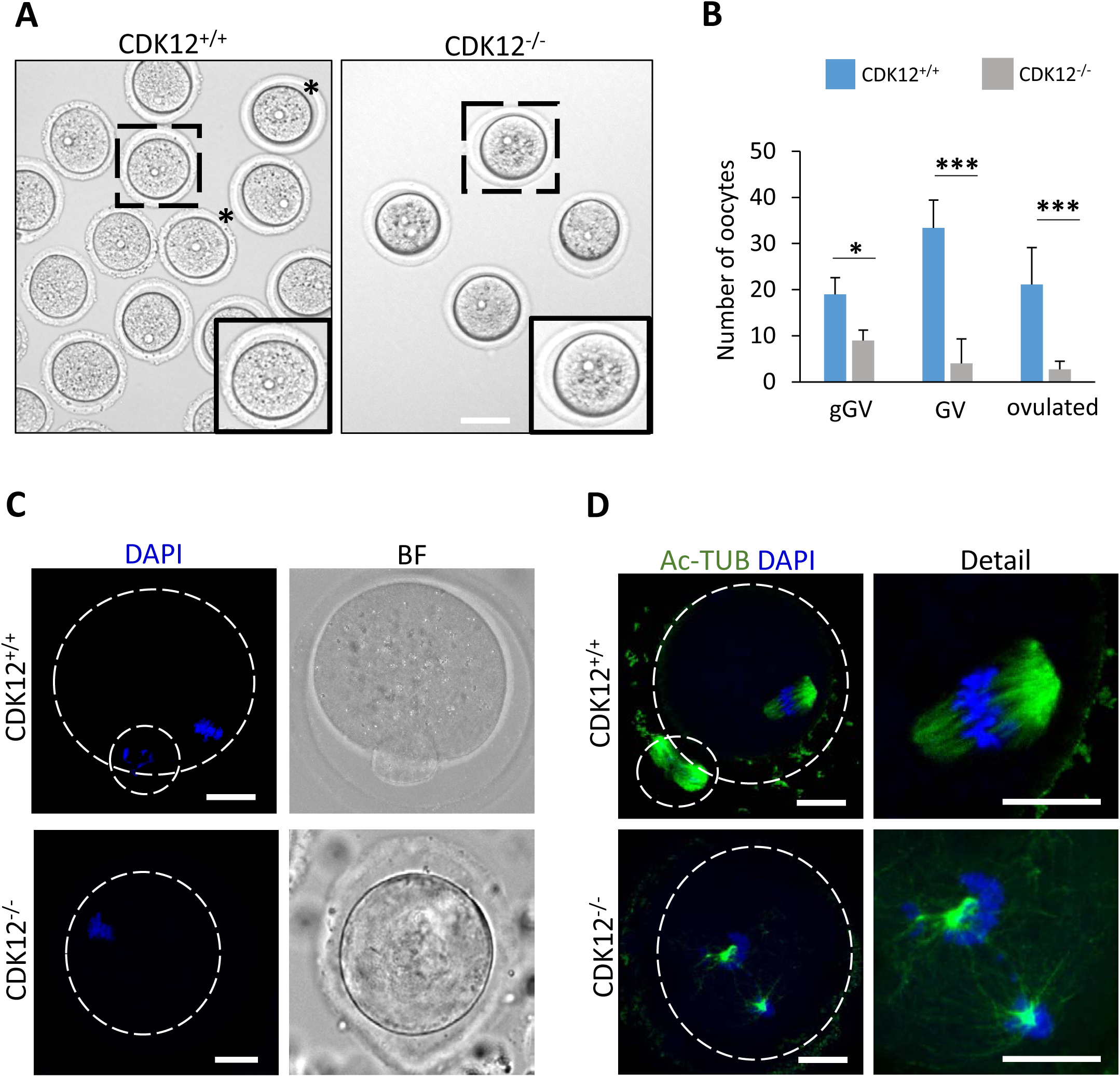
CDK12 is Essential for oocyte development and maturation. **A)** Representative images of morphology of CDK12^+/+^ and CDK12^-/-^ oocytes. Dashed squares show details of oocytes, asterisks show fully grown oocytes; scale bars 60 µm. **B)** Quantification of growing (gGV), fully-grown (GV) and ovulated (post hCG) oocytes isolated from ovaries. Data from five independent biological replicates. Data are presented as mean ± SE; Student’s t-test: *p < 0.05; ***p < 0.001. **C)** Morphology of CDK12^+/+^ and CDK12^-/-^ ovulated oocytes. Data from three biological replicates and n≤ 30 cells per group. DAPI (blue), bright field (BF); the dashed line shows the cell cortex; scale bars 20 µm. **D)** Representative images of spindle morphology labelled with acetylated α-tubulin (Ac-TUB, green) in ovulated CDK12^+/+^ and CDK12^-/-^ oocytes. Data from three biological replicates and n≤ 27 per group. Details show the enlargement of the spindle area. DAPI (blue); the dashed line shows the cell cortex; scale bars 20 µm.

In summary, the absence of CDK12 in the oocyte leads to an interruption of development to fully grown GV stage and thus to a termination of folliculogenesis, resulting in female infertility.

### The absence of CDK12 suppresses transcriptional activity in developing oocytes

Our results suggest that the main cause of infertility is impaired development of the growing oocytes. Previous reports indicated a close link between CDK12 and the regulation of transcription via phosphorylation of RNA Polymerase II (POLII) [10]. We hypothesized that this regulatory function of CDK12 is critical for proper gene expression during oocyte growth. First, we found that CDK12 is predominantly localized in the oocyte nucleus (**Supporting Information Figure 1C**). In addition, proximity ligation shows that CDK12 is localized together with its binding partner cyclin K (CCNK) in the nucleus of the mouse and human oocyte (**Supporting Information 3**). To assess overall transcription, we next labeled newly synthesized RNA with 5-ethynyluridine (EU) in growing oocytes. CDK12^-/-^ oocytes exhibited a 69% decrease in EU staining compared to CDK12^+/+^ oocytes, indicating a significant decrease in global transcription (**Figure 4 A, B**). In addition, the active form of RNA Polymerase II (Ser2), a marker for transcription elongation, was reduced by 39% in CDK12^-/-^ oocytes (**Figure 4C, D**). Importantly, there were no differences in PolII mRNA and protein levels between groups (**Figure 5A, B**).

**Figure 4:**
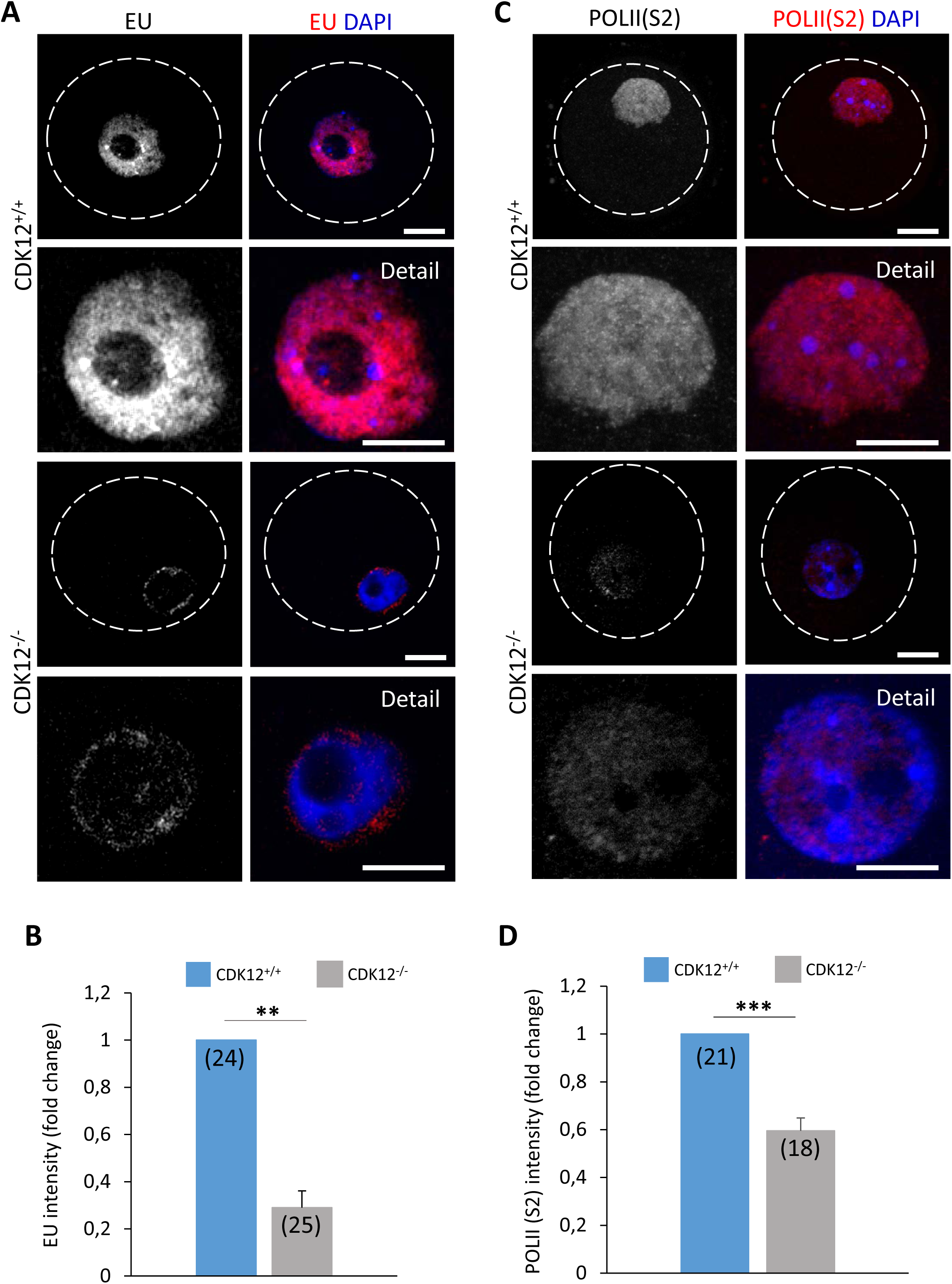
The absence of CDK12 supress transcriptional activity in developing oocytes. **A)** Detection of transcriptional activity with 5-Ethynyl Uridine (EU; gray and red) in CDK12^+/+^ and CDK12^-/-^ oocytes. Data from three independent biological replicates. Details show a magnification of the nucleus; DAPI (blue); the dashed line depicts the cell cortex; scale bar 20 µm. **B)** Quantification of EU fluorescence in the nucleus from (**A**). The values from CDK12^+/+^ were set as 1. The number of cells is shown in parentheses. Data are presented as mean ± SE; Student’s *t*-test: **p < 0.01. **C)** Detection of phosphorylation of Polymerase II at Serine 2 (POLII(S2); gray and red) by immunocytochemistry. Data from three independent experiments. The details show an enlargement of the nuclear region. DAPI (blue); the dashed line depicts the cell cortex; scale bar 20 µm. **D)** Quantification of POLII (S2) fluorescence intensity in the nucleus of CDK12^+/+^ and CDK12^-/-^ oocytes from the (**C**) experiment. The values from CDK12^+/+^ were set as 1. The number of cells is shown in parentheses. Data are presented as mean ± SE; Student’s *t*-test: ***p < 0.001.

**Figure 5:**
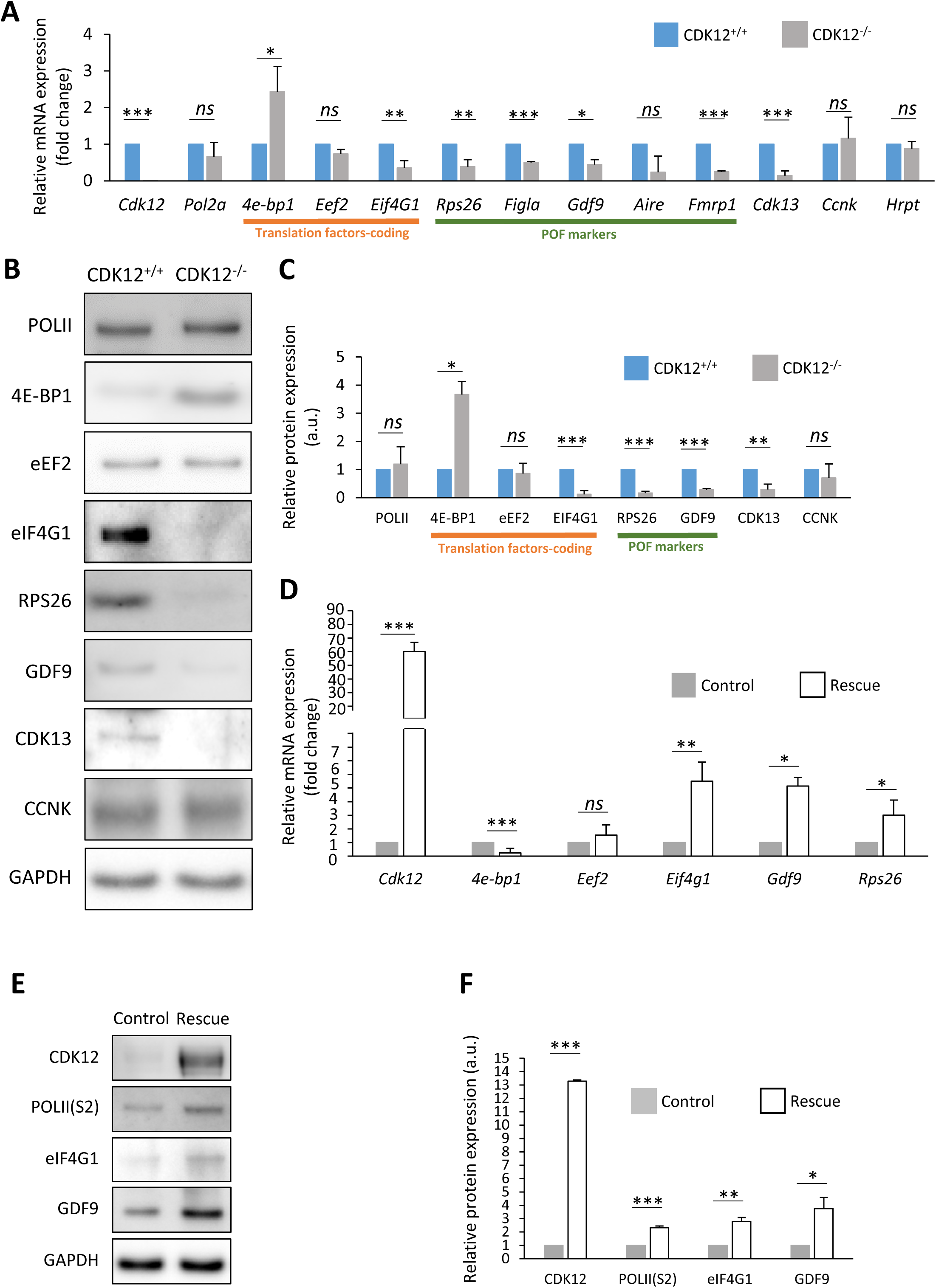
The absence of CDK12 influences the expression of a subset of mRNAs in the oocyte. **A)** Quantitative RT-PCR analysis of selected mRNAs. RNAs coding for translational factors (underlined in orange) and markers for premature ovarian failure markers (underlined in green). Data normalized to *Hrpt* mRNA. Data from three independent biological replicates. Values from CDK12^+/+^ oocytes were set as 1. Data are presented as mean ± SD; Student’s *t*-test: *ns*, non-significant; *p < 0.05, **p<0.01; ***p < 0.001. **B)** Western blot analysis of candidate proteins selected from (**A**). Values from CDK12^+/+^ oocytes were set as 1. GAPDH was used as a loading control. Data from at least three independent biological replicates. For the phosphorylation of 4E-BP1, see Fig. 3. **C)** Quantification of candidate proteins from (**B**). Data are presented as mean ± SE; Student’s *t*-test: *ns*, non-significant; *p < 0.05; ***p < 0.001. **D)** Quantitative RT-PCR analysis of the expression of selected mRNAs in control (microinjection of *H2b-Gfp* RNA into CDK12^-/-^ oocytes; gray) and in CDK12^-/-^ oocytes microinjected with *Cdk12 m*RNA (rescue; white). Data from three independent biological replicates. Values were normalized to the number of oocytes per sample. Data are presented as mean ± SD; Student’s *t*-test: *ns*, non-significant; *p < 0.05. **E)** Western blot analysis of selected proteins after CDK12 overexpression. GAPDH was used as a loading control. Data from three biological replicates. **F)** Quantification of candidate proteins from (**E**). Data are presented as mean ± SE; Student’s *t*-test: *ns*, non-significant; *p < 0.05; **p<0.01; ***p < 0.001.

These results suggest that the absence of CDK12 leads to an abnormal transcriptional program that impairs the formation of the oocyte mRNA reserve and consequently prevents the development to fully grown GV oocytes and the depletion of the ovarian reserve.

### The absence of CDK12 influences the expression of a subset of mRNAs in the oocyte

It is well known that oocyte development depends on proper gene expression. Given the significant reduction in transcription observed in CDK12^-/-^ oocytes (**Figure 4**), we examined the expression of specific developmentally relevant classes of mRNAs encoding translational factors [7] (4E-BP1, eEF2, eIF4G1) and markers associated with premature ovarian failure [15–18] a phenotype observed in (**Figure 2**) (POF; RPS26, FIGLA, GDF9, AIRE, FMRP1). Interestingly, qPCR data showed a significant 3.5-fold increase in mRNA of the translational repressor *4e-bp1* in CDK12^-/-^ oocytes (**Figure 5A**). However, the mRNAs encoding the translation elongation factor EEF2 were similar, and the mRNA encoding the translation initiation factor eIF4G1 was significantly decreased in CDK12^-/-^ oocytes (**Figure 5A**). In addition, all analyzed mRNAs encoding POF markers [16,18,19] were significantly reduced in CDK12^-/-^oocytes (**Figure 5A**). In addition, we analyzed the expression of mRNAs encoding the homologous kinase CDK13 and the CDK12 binding partner CCNK (**Supporting Information 3**). The mRNAs of CDK12 and CDK13 were significantly decreased in CDK12^-/-^ oocytes, whereas the mRNAs of CCNK, POLII and HRPT were equally expressed in both groups (**Figure 5A**). Next, we analyzed the protein expression of selected candidate genes that had previously been examined by PCR (**Figure 5A**). Immunoblotting analysis showed a positive correlation of the expression of POLII, 4E- BP1, eEF2, eIF4G1, RPS26, GDF9, CDK13 and CCNK with the corresponding mRNAs (**Figure 5B, C**). To analyze the direct effect of CDK12 on the expression of candidate genes, we microinjected RNA encoding mouse CDK12 into CDK12^-/-^ oocytes. We restored the expression of CDK12 in the CDK12^-/-^ oocytes (**Figure 5D-F**) and simultaneously observed decreased expression of the mRNA of the translational repressor *4e-bp1*. CDK12 expression increased the mRNAs of the *Eif4g1*, *Gdf9* and *Rps26* (**Figure 5D**). Importantly, the presence of CDK12 in CDK12^-/-^ oocytes promoted the increase in POLII phosphorylation at serine 2 as well as the expression of eIF4G1 and the POF marker GDF9 (**Figure 5E, F**).

In summary, the absence of CDK12 affects the expression of some selected mRNAs related to translation and the premature ovarian failure phenotype. Furthermore, the addition of exogenous CDK12 to CDK12^-/-^ oocytes enhances POLII activity, leading to a reorganization of the expression of translation factors and markers for POF.

### Aberrant transcriptome of CDK12^-/-^ oocytes represses global translation and enhances expression of the translational repressor 4E-BP1

Considering that the absence of CDK12 promotes an aberrant maternal transcriptome and in particular translational factors in the developing oocyte (**Figure 5A - C**), we investigated how protein synthesis was affected in CDK12^-/-^ oocytes. First, a ^35^S-methionine incorporation assay, showed that CDK12^-/-^ oocytes exhibited a 23 % reduction in global protein synthesis compared to WT oocytes (**Figure 6A, B**). Considering the reported effect of CDK12 on RNA polyadenylation[20,21] and the impaired transcriptome/proteome (**Figure 5A-D**) in CDK12^-/-^ oocytes, the polyadenylation directly correlates with the rate of protein synthesis of mRNA. We examined global polyadenylation by RNA FISH, which showed no difference between CDK12^+/+^ and CDK12^-/-^ oocytes (**Figure 6C and Supporting Information 4**). In connection to overexpression of the translational repressor 4E-BP1 and reduced translation in CDK12^-/-^ oocytes (**Figure 5A, B**), we analyzed polyadenylation of the mRNA encoding 4E-BP1. The PAT assay showed a visible poly(A) shift of the 3’UTR tail of *4e-bp1* in CDK12^-/-^ oocytes, while the poly(A) tail remained unchanged in *Cnot7 mRNA,* which is translated after completion of meiosis I [22] (**Figure 6D, E**).

**Figure 6:**
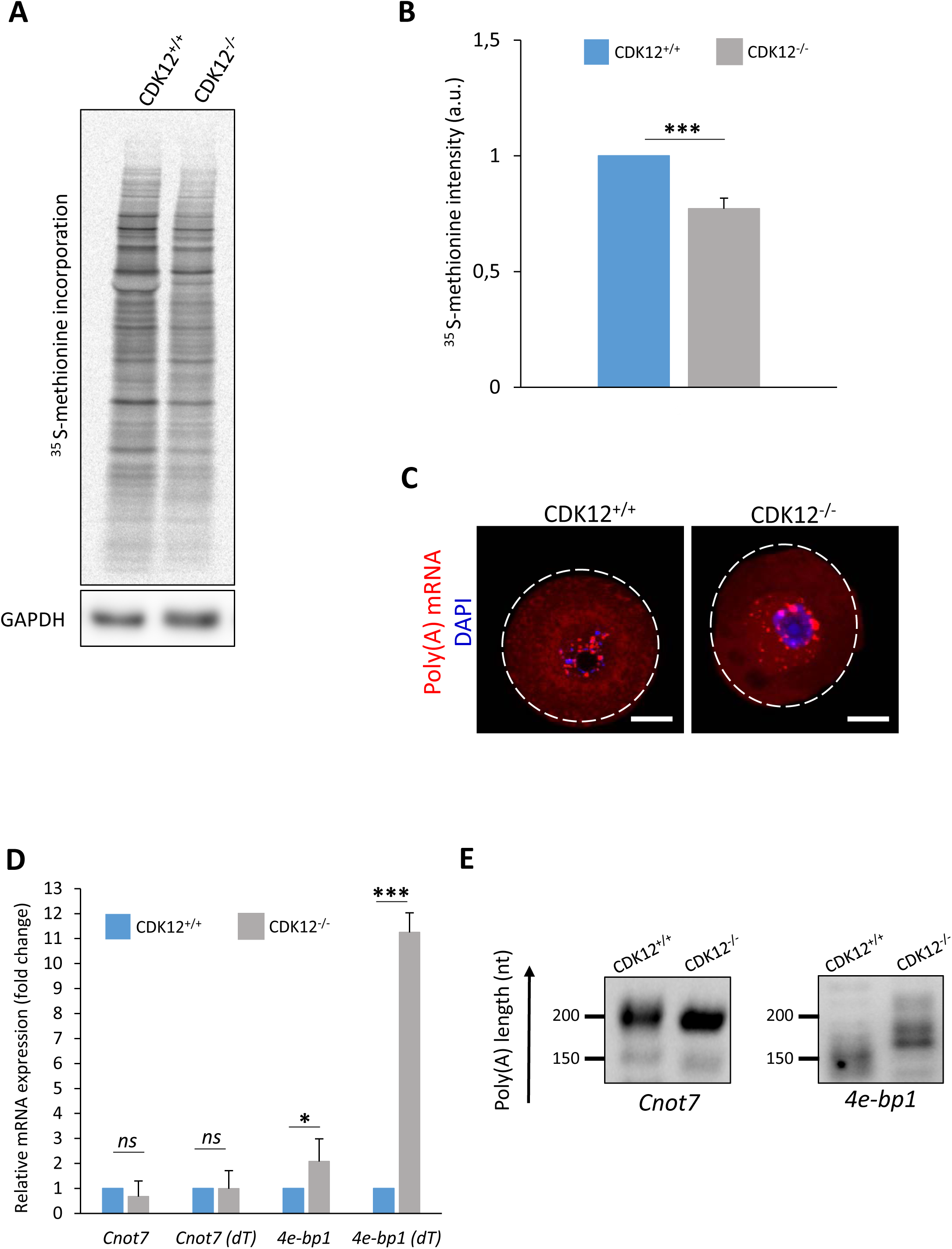
The absence of CDK12 specifically affects the stability of the mRNA encoding the translational repressor 4E-BP1. **A)** Analysis of global *de novo* proteosynthesis by incorporation of ^35^S-methionine in CDK12^+/+^ and CDK12^-/-^ oocytes. Data from four independent biological replicates. GAPDH was used as a loading control. **B)** Quantification of ^35^S-methionine incorporation from (**A**). Values from CDK12^+/+^ oocytes were set as 1. Data from four independent experiments. Data presented as mean ± SD; Student’s *t*-test: ***p < 0.001. **C)** Representative confocal images of poly(A) mRNA in CDK12^+/+^ and CDK12^-/-^ oocytes. Data from three biological replicates. Poly(A) mRNA (red); DAPI (blue); the dashed line depicts the cell cortex; scale bars 20 µm. For the quantification of fluorescence intensity, see SI Fig. 4. **D)** Quantitative RT-PCR analysis of cDNA synthetized with hexamers or oligo(dT) primers in CDK12^+/+^ and CDK12^-/-^ oocytes. Data from three biological replicates. Data are normalized to the number of oocytes. Data are presented as mean ± SD; Student’s t-test: *ns*, non-significant; *p < 0.05; ***p < 0.001. **E)** Polyadenylation test (PAT) to detect the poly(A) tail length of *4e-bp1* in CDK12^+/+^ and CDK12^-/-^ oocytes. Translationally inactive *Cnot7* mRNA was used as negative control. Data from three independent biological replicates. The length of the poly(A)tail is described on the left (nt).

In summary, we document that the absence of CDK12 leads to the stabilization of specific mRNAs, in particular *4e-bp1*, which is in the non-phosphorylated state (**Supporting Information 5**), thereby acting as a translational repressor and leading to reduced protein synthesis.

Our results clearly show that the absence of CDK12 leads to maternal infertility via the disruption of oocyte growth by a mechanism that impairs maternal transcriptome and translation (Graphical abstract). In addition, discontinued oocyte development leads to the failure of folliculogenesis, which is similar to the afollicular form of premature ovarian failure that also occurs in humans [23].

## Discussion

We have summarized here the biological function of CDK12 with a focus on female reproduction. Interestingly, the absence of CDK12 in the oocyte results in the absence of fully developed oocytes, leading to complete female sterility. With Cre, CDK12 is downregulated under the ZP3 promoter, which is activated in the growing 20 µm oocyte and reaches its maximum in the 50 µm oocyte [24]. This approach allows us to investigate the role of CDK12 in the transcriptionally active phase of oocyte development, which influences the early phase of oocyte growth and thus folliculogenesis. Although ovulation is promoted in the absence of CDK12, it is only sparse and oocytes do not mature to a stage that can be fertilized and at least promote subfertility.

The CDK12/CCNK complex has been shown to be required for the promotion of transcription elongation by RNA polymerase II and the phosphorylation of its carboxyl-terminal domain [25]. CDK12 has sequences for nuclear localization[26]. In the oocyte, CDK12/CCNK complexes are abundant in the nucleus, highlighting their biological function in transcription [8,27]. CDK12 and CDK13 exhibit a high degree of homology and exhibit partial redundancy in their function in polymerase II-mediated transcription [8,28]. In cancer cell lines, the genetic deletion or chemical inhibition of CDK12 kinase activity has virtually no effect on CDK13 protein levels, and never leads to such a dramatic downregulation as in growing oocytes [27,29]. It is therefore difficult to decide whether the 69% decrease in global transcription is due to the absence of CDK12 or CDK13. In cancer cell lines, inhibition of the kinase activity of both kinases by several functionally different inhibitors affected transcription efficiency by 10 - 20% [5,28]. One can speculate that the role of CDK12 and CDK13 in regulating transcription in cancer cell lines is different from that in primary cells such as oocytes. Not surprisingly, the decrease in global transcription was accompanied by a 39% decrease in the Ser2 phosphorylation labelling of elongating Pol II. Since we could still detect Ser2 signaling in the nucleus, it is very likely that CDK9 is the kinase responsible for the remaining Pol II(Ser2) signal, as CDK9 is active in the GVs of porcine oocytes, and their inhibition by CDK9 resulted in developmental arrest [30]. Furthermore, the chemical inhibition of CDK12, CDK13 or both together affected up to 15 % of all genes, clearly indicating that the transcriptional program in growing oocytes is mainly the transcription of specialized and developmentally relevant genes [28]. It is likely that the downregulation of transcription in the cKO oocytes only affects the transcription of specialized genes. Although we did not perform a global transcriptome analysis, we clearly found decreased mRNA for POF markers and translation factors with the sole exception of the *4e-bp1* mRNA. Suggesting that the absence of CDK12 **i**) promotes the transcription of specific genes, **ii**) impairs a specific transcriptional program during the activation of the ZP3 promoter in the oocyte, and/or **iii**) indirectly affects transcription by altering the expression of factors that inhibit the transcription process. Our results show that the absence of CDK12 affects the transcriptome of the growing oocyte by interfering with the activity of Pol II, which in turn affects translation. Previous reports showed that the yeast ortholog of CDK12, Ctk1 [8,31] stimulates mRNA translation globally [12,32]. Similarly, we found an additional role of CDK12 that involves the stabilization of specific mRNA with increased polyadenylation, thereby promoting their translation. Indeed, CDK12 has been identified as a nuclear co-transcriptional polyadenylation factor that either ensures the processing of the 3’-end through a cleavage and polyadenylation mechanism, or blocks the premature cleavage and polyadenylation (PCPA) of hnRNA [21,33]. Apart from the effect of CDK12 on the polyadenylation of 4E-BP1 mRNA, the deletion of CDK12 resulted in a moderate increase in the protein level of the 5’-cap-binding mRNA repressor 4E-BP1 in CDK12^-/-^ oocytes. Our previously published results showed that expression of the inactive translational repressor 4E-BP1 leads to aberrant proteosynthesis in fully developed, transcriptionally silent oocyte [34]. Most mRNAs accumulated during oocyte growth are translationally repressed and are translated later when transcription is repressed [35,36]. Therefore, 4E-BP1 may play a key role in this process, and the overexpression of 4E-BP1 observed in CDK12^-/-^ oocytes suppresses the translation of already abnormally expressed mRNAs, the translation of which promotes oocyte growth. We previously, reported that inactivation of 4E-BP1 occurs at later stages of oocyte development, after entry into the M phase [34], and similarly, CDK12 phosphorylates/inactivates 4E-BP1 and promotes the translation of specific mRNA of factors involved in mitotic spindle regulation and chromosome segregation [7]. Importantly, the translational repressor 4E-BP1 is inactivated by phosphorylation during the metaphase transition in cells and oocytes [7,34]. The unphosphorylated form of 4E-BP1 is therefore present in growing GV oocytes. Nevertheless, we still do not know whether and how the mRNA stabilization and translation of 4E-BP1 is directly or indirectly influenced by CDK12. In addition, the expression of the translation initiation factor eIF4G1 and aberrant transcriptome in CDK12^-/-^ oocytes, likely contributing to reduced global translation.

Among the downregulated transcripts and proteins, premature ovarian failure (POF) markers such as *Rps26*, *Figla*, *Gdf9*, *Aire* and *Fmrp1* [16,17,37–39] were detected, so the absence of CDK12 clearly resembles a POF phenotyp [40–42]. POF is a clinical disorder characterized by hypogonadism and amenorrhea, and affects 1– 3% of women under 40 years of age [43–46]. The overall prevalence of familial POF ranges from 4% to 31% [45]. The appearance of the ovaries varies from the complete depletion of follicles to the presence of a variable population of follicles that fail to develop [43,47]. GDF9 and RPS26 have been described as important for oocyte growth and the recruitment of a primordial follicle into the pool of growing follicles [17,48]. GDF9, a protein secreted from the oocyte into the follicle, influences the proliferation, differentiation, steroid hormone synthesis, apoptosis and cumulus expansion of granulosa cells [49]. GDF9 levels increase dramatically with oocyte growth during preantral folliculogenesis and remain high until ovulation [48] in oocytes of various mammalian species, including humans [50] (sheep [51], bovines [51] and rats [52], indicating the universality of the role of GDF9. In addition, CDK12 has been shown to be associated with follicular atresia (reduction in the number of ovulating follicles) and early menopause in humans [40,42].

Taken together, our results clearly show that the absence of CDK12 leads to an abnormal transcriptome and translatome of the growing oocyte, which in turn results in the absence of a fully mature oocyte and leads to female infertility. In addition, we provide insight into the etiology of premature ovarian failure, in which CDK12 may also play an important role in humans, warranting further investigation.

## Methods

### Oocyte isolation and cultivation

Experimental genotypes were designated as wild-type (WT; CDK12^+/+^; Cdk12^tm1c+/+^ Zp3-Cre^+/+^) and homozygotes (cKO; CDK12^-/-^; Cdk12^tm1c-/-^ Zp3^+/+^). Oocytes were obtained from C57BL/6J mice that were at least 5 weeks of age. Females were stimulated with 5 IU of pregnant mare serum gonadotropin (PMSG; Folligon; Merck Animal Health) per mouse 46 hours prior to oocyte isolation. To obtain MII, the mice were primed with 5 IU human chorionic gonadotropin (hCG, Pregnyl, N.V. Organon). Oocytes were isolated in transfer medium (TM) supplemented with 100 µM 3-isobutyl-1- methylxanthine (IBMX, Sigma Aldrich) to prevent the spontaneous resumption of meiosis. Selected oocytes were denuded and cultured in M16 medium (Millipore) without IBMX at 37 °C, 5% CO_2_ for 0 h (GV) or 12 h (MII) [53]. All animal experiments were performed in accordance with the guidelines and protocols approved for the Laboratory of Biochemistry and Molecular Biology of Germ Cells at the Institute of Animal Physiology and Genetics in the Czech Republic. All animal experiments were conducted in accordance with Act No. 246/1992 on the Protection of Animals from Cruelty, issued by the Ministry of Agriculture under the number 67756/2020MZE-18134. Human oocytes not used for human reproduction were obtained from the Institute for the Care of Mother and Child in Prague. The project was approved and accredited by the Ethics Committee of the Institute for the Care of Mother and Child (#1/3/4/2022; NU-23-07-0005). All patients gave informed consent to the use of their immature oocyte(s) this study.

### Animals

Cdk12^fx/fx^ mice on a C57BL/6J genomic background were provided by Jiri Kohoutek. Cdk12^fx/fx^ mice were crossed with Zp3-Cre mice to generate the oocyte-specific Cdk12 knockout (CDK12^-/-^). The target construct contains loxP sites in the 3 and 4^th^ exon (8 nucleotides in the defective 4th exon and a stop codon inserted into the open reading frame). The primers used for PCR to genotype Cdk12^fx/fx^ were primer mCDK12 Flx-F–CTTCAGACAGTGTCAGACCACCTGGAGAAGC; primer mCDK12 Y3-R–CCTCTGACCTCCCAATGTGTGCATGACAC; F-ZP3-GGTGGAGAATGTTAATC and R-ZP3-TATTCGGATCATCAGCTA. All experiments were performed according to the guidelines and with the approval of Institutional Animal Care. Pairs of 8-week-old female mice of genotypes CDK12^+/+^, CDK12^+/-^ and CDK12^-/-^ were continuously mated with proven CDK12^+/+^ males to test the fertility of the females. The number of pups was recorded over a period of 8 months.

### Histology of ovaries

Ovaries were fixed in 4% PFA solution (Sigma) for 48 h, then placed in 70% ethanol solution and subsequently placed in labeled histological cassettes. Samples were processed using an automated tissue processor (Leica ASP 6025, Leica Microsystems, Germany) and embedded in paraffin blocks using a Leica EG 1150H paraffin embedding station (Leica Microsystems, Germany). Sections of 3 μm were cut with a microtome (Leica RM2255, Leica Microsystems, Germany) on standard glass slides (Waldemar Knittel, GmbH, Germany), every 10th section was collected, 3-12 sections were collected per slide. Slides were stained with hematoxylin–eosin and mounted using a Leica ST5020 automated stainer in combination with a Leica CV5030 mounting frame. The number of follicles was quantified using a stereomicroscope (Zeiss Stemi 2000, Germany). The analysis of ovarian follicles was performed by two different investigators.

### Oocyte microinjection

Experimental oocytes Cdk12^-/-^ were injected with an *in vitro* prepared *Cdk12* and *H2b-Gfp* RNAs diluted to a final concentration of 20 ng/µl, the control samples were injected with *H2b-Gfp* mRNA alone. Subsequently, the injected oocytes were incubated in IBMX at 37 °C and 5% CO_2_ for 18 h. The oocytes were washed in PVA/PBS and frozen at − 80°C.

#### Measurement of Overall Protein Synthesis

To measure total protein synthesis, 50mCi of ^35^S-methionine (Perkin Elmer) was added to methionine-free culture medium, for 1 h and then oocytes were lysed in SDS-buffer and subjected to SDS–polyacrylamide gel electrophoresis. The labelled proteins were visualized by autoradiography with BasReader (FujiFilm). GAPDH was used as a loading control.

### 5-ethynyl uridine transcription assay

The 5-EU was added to the M16 medium with IBMX and incubated overnight with growing GV oocytes. Oocytes were then fixed in 4% paraformaldehyde/PBS for 15 min, permeabilized with 0.1% Triton X-100/PBS for 10 min at room temperature, and incubated for 1 h at room temperature in the dark with the Click-iT reaction cocktail (according to the manufacturer’s instructions using Alexa A555 azide (ThermoFisher, A20012) and a commercial kit (Click-iT, ThermoFisher, C10276). After incubation, oocytes were washed once with PBS and mounted on slides using DAPI in the presence of the anti-fade reagent Vectashield (H-1500, Vector laboratories). Images were captured using a confocal laser scanning microscope (Leica SP5, Leica Microsystems, Wetzlar, Germany). Images were quantified and compiled using FIJI software (version 1.8.0_172).

### RNA isolation and RT-PCR

TRIzol reagent (Invitrogen) was used for RNA extraction according to the manufacturer’s instructions. Reverse transcription was performed with a qPCRBIO cDNA Synthesis Kit (PCR Biosystems). qPCR was then carried out using QuantStudio 3 (Applied Biosystems) and Luna® Universal qPCR Master Mix (New England BioLabs) according to the manufacturer’s protocols with an annealing temperature of 60°C. The primers are listed in Table 1.

**Table 1.**
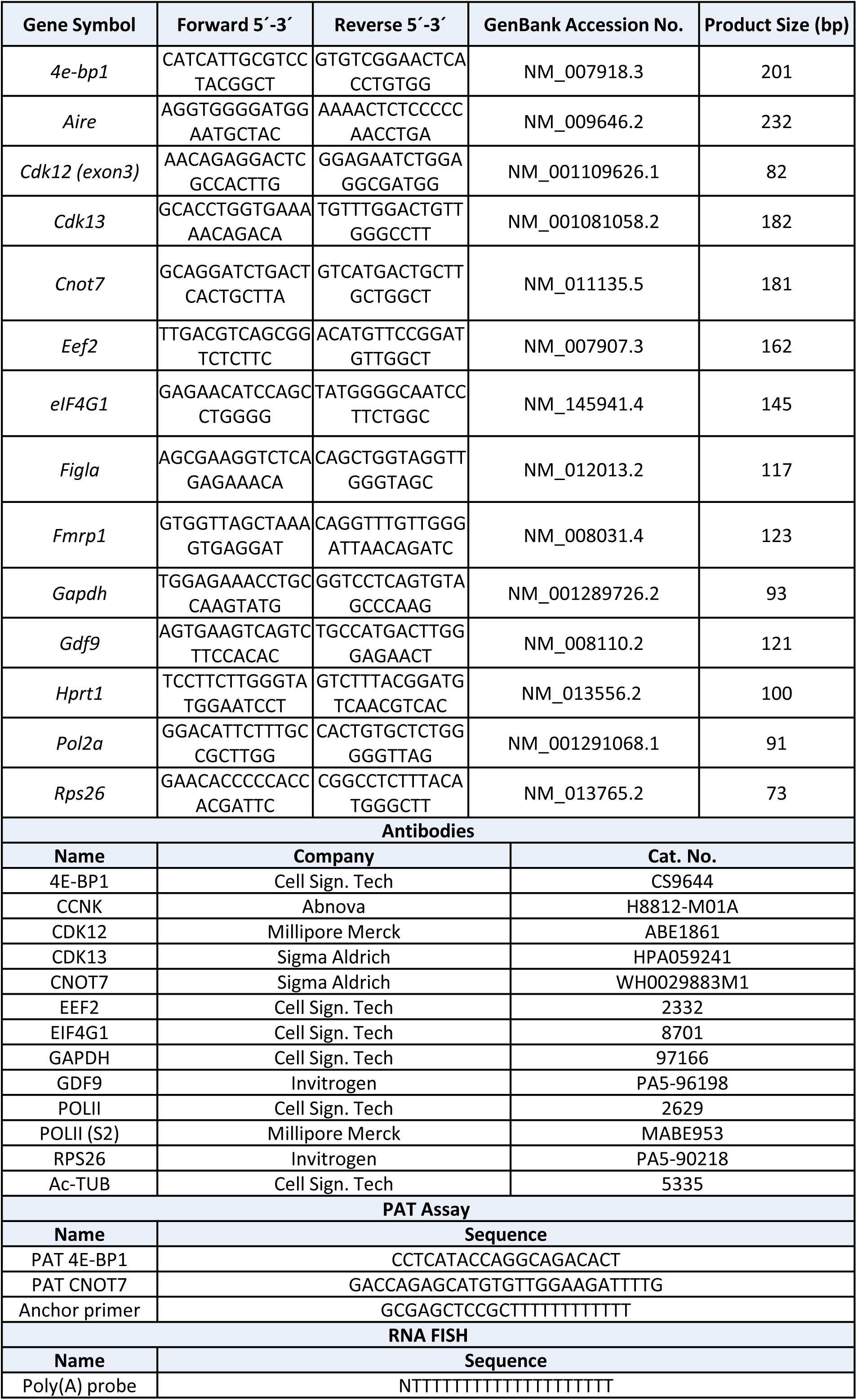

### Poly-A-tail length assay (PAT)

To obtain the total length of the poly(A) tail of each transcript, total RNA was extracted using the phenol-chloroform method according to the laboratory protocol. Elution was performed in 10 µl of water per sample. The isolated RNA was incubated with 1 µl of 20mM oligo-dT annealing per sample for 5 min at 65°C. Ligation was then carried out for 30 min at 42°C. The ligation mix was prepared from the following components: T4 ligase (1 µl), Superscript IV 5x buffer (5 µl), 20U/µl RNAse inhibitor (1 µl), 10mM dNTP (1 µl), 10 mM ATP (1 µl), 1M MgCl2 (0.1 µl), 0.1M DTT (2 µl), RNAse- free water (2 µl). The cDNA synthesis was performed by adding 1 µl Superscript II Reverse Transcriptase with the following setup: 45 min at 45°C, 10 min at 80°C, hold at 4°C. The prepared cDNA was subjected to PCR with gene-specific forward primers and anchoring reverse primer (Table 1). PCR was performed with PPP Master Mix (Top-Bio) under the following conditions: 1 min at 95°C, 35x (30 s at 95°C, 20 s at 55°C, 45 s at 72°C). PCR products were analyzed on a 1.5% agarose gel stained with GelRed (41003, Biotinum) and run at 90 V for 45 min. The gels were detected with an Azure 600 Imager (Azure Biosystems).

### RNA FISH of poly(A) mRNA

Oocytes were fixed in 4% paraformaldehyde for 15 min and permeabilized by protease III treatment (Biosearch Technologies). Samples were washed in wash buffer A (Biosearch Technologies) and incubated overnight at 42°C in hybridization buffer (Biosearch Technologies) containing 75 nM oligo-d(T) probe with CalFluorRed635 (Biotech Generi). Samples were washed twice in wash buffer A and twice in 2x SSC (Sigma Aldrich). Samples were embedded in VectaShield medium with DAPI (H-1500, Vector Laboratories). A confocal laser scanning microscope was used for imaging (Leica SP5, Leica Microsystems, Wetzlar, Germany). The quantification of fluorescence intensity between CDK12^+/+^ and CDK12^-/-^ oocytes was performed using FIJI software (version 1.8.0_172).

### Immunofluorescence

Fixed oocytes (15 min in 4% PFA, Sigma Aldrich) were permeabilized for 10 min in 0.1% Triton X-100, washed in PBS with polyvinyl alcohol (PVA, Sigma Aldrich), and incubated overnight at 4 °C with primary antibodies (Table 1) diluted in PVA/PBS. Immunofluorescence analysis was performed according to a published protocol [54]. Image quantification and compilation was performed using FIJI software (version 1.8.0_172).

#### In situ proximity ligation assay (PLA)

The proximity ligation assay was performed according to the instructions of Naveni Triflex ( Navinci). Oocytes were fixed in 4% PFA for 15 min and permeabilized with 0.5% TritonX/PBS for 10 min. Oocytes were washed with TBS-T solution and then transferred into primary antibodies (CCNK and CDK12, Table 1) 1.5: 100 dilution 1TF overnight at 4°C. The oocytes were washed in TBS-T for 15 min. The oocytes were incubated with Navenibody MTF and Navenibody RTF 1:10 dilution 2 for 1 h at 37°C on a hot plate. The oocytes were washed every 15 min with TBS-T solution. 40 µl of amplification reaction 1 was mixed according to the manufacturer’s instructions, then added to the oocytes and incubated at 37°C for 30 min. It was then washed with TBS-T solution for 5 min. Reaction 2 was mixed according to the manufacturer’s instructions. The oocytes were incubated in 40 µl of reaction 2, in a dish protected from light, for 1 h at 37°C. Samples were washed for 5 min and then mounted on a concave-bottomed slide (cat. # 1216492, Marienfeld,) using Vectashield mounting medium with DAPI (H-1500, Vector Laboratories). A confocal laser scanning microscope was used for the images (Stellaris8).

#### Western Blotting

Oocyte lysates were analyzed on a 4–12% gradient acrylamide gel. Samples were transferred to a polyvinylidene fluoride membrane (Immobilon P; Merckmillipore) using a blotting system (Biometra GmbH) at 5 mA/cm2 for 25 min. The membranes were blocked for 1 h at room temperature and then incubated overnight at 4 ᵒC with the primary antibodies listed in Table 1. Membranes were incubated with secondary antibodies for 1 h at room temperature. Proteins were visualised by chemiluminescence using ECL (Amersham) and imaged in an Azure 600 Imager (Azure Biosystems), and the acquired signals were quantified using ImageJ (http://rsbweb.nih.gov/ij/). To detect the phosphorylation shift, oocytes were dissolved in 20 µl of 1x NEB buffer containing 800 U of LPP enzyme (P0753, New England BioLabs) and incubated at 30°C for 1 h.

#### Statistical analysis and data visualization

GraphPad Prism 8.3 was used for the statistical analysis. The statistical analysis included Student’s t-tests to determine statistical significance between groups (labelled with an asterisk). *p < 0.05; **p < 0.01 and ***p < 0.001. Mean and standard error values were calculated in MS Excel (Microsoft).

## Acknowledgments

We wish to thank J. Supolikova and M. Hancova for technical assistance. We also thank to A. Jindrova, R. Iyyappan and K. Kakavand for critical reading of the manuscript. The authors used the services of the Czech Center for Phenogenomics at IMG supported by the Czech Academy of Sciences RVO 68378050 and the project LM2023036. This work was supported by GACR 22-27301S, EXCELLENCE [CZ.02.1.01/0.0/0.0/15_003/0000460 OP RDE], Institutional Research Concept RVO67985904, and V. Sedmikova by IGA from IAPG 2022.

## Conflict of interest statement

None declared.

## Author Contributions

**D.J.:** Supervision, Conceptualization, Investigation, Validation, Methodology, Writing – original draft, review & editing. Project administration. **V.S.:** Validation, Methodology, Investigation. **F.B.:** Validation, Methodology, Investigation, review & editing. **D.A**.: Validation, Methodology, Investigation. **M.D.:** Investigation. **M.K.:** Funding acquisition, editing. **J.R.:** Methodology. **J.R.:** Methodology. **J.K**.: Writing – original draft, review & editing. **A.S.:** Conceptualization, Supervision, Funding acquisition, Writing – original draft, review & editing.

---

**SI Fig. 1:**
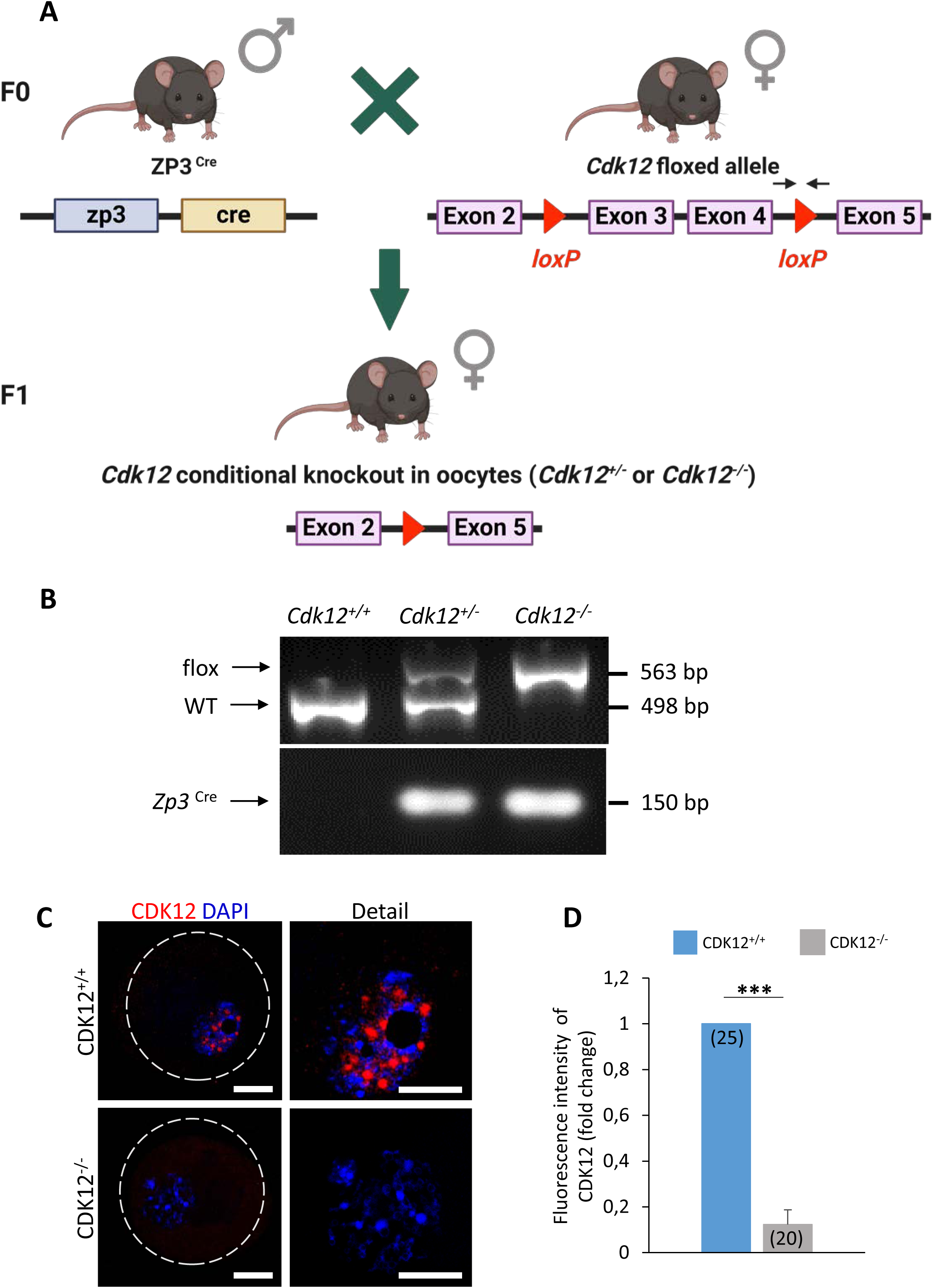
The generation of mice with conditional knockout of CDK12 in the oocyte. **A)** Scheme of CDK12 conditionally deleted via the promoter of Zona Pellucida 3 (ZP3)-driven Cre-Lox recombinase in the oocyte. Black arrows indicate the position of the primers for genotyping. **B)** PCR for genotyping genomic DNA isolated from different genotypes. Genotyping of progeny for the presence of CDK12(^tm1c)(flox^) and Zp3^-Cre^. The wild-type allele produces a 498-bp product, while the *flox* allele produces a 563-bp product. Zp3^-Cre^ generates a 150 bp product. **C)** Immunocytochemical analysis of CDK12 expression and localization in oocytes. Data from three independent biological replicates. CDK12 (red); DAPI (blue); dashed line depicts cell cortex; scale bar 20 µm. **D)** Quantification of CDK12 fluorescence intensity in CDK12^+/+^ and CDK12^-/-^ oocytes from (**C**). The number of cells is shown in parentheses. Values from CDK12^+/+^ oocytes were set as 1. Data are presented as mean ± SE; Student’s t-test: ***p < 0.001.

**SI Fig. 2:**
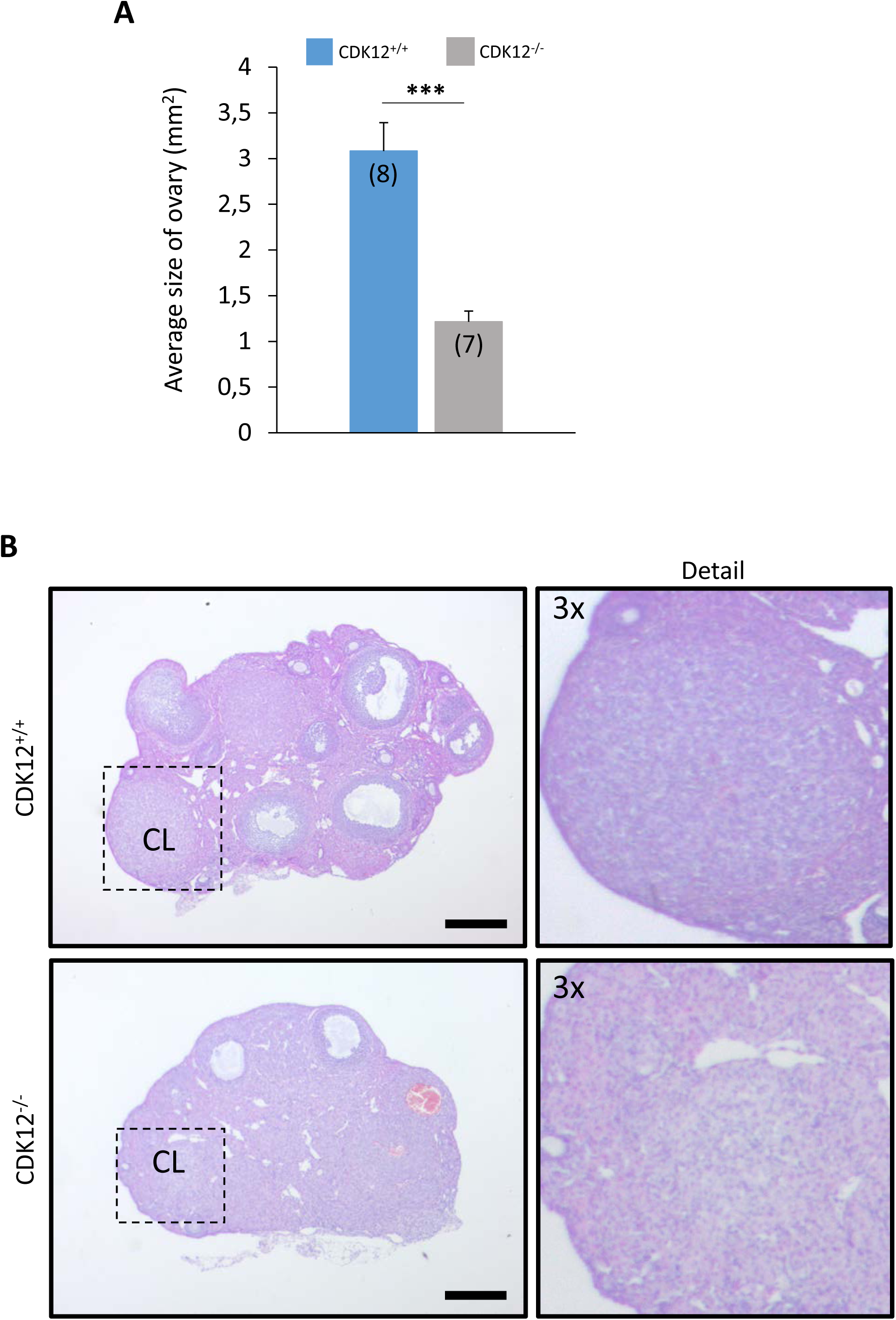
The absence of CDK12 in oocytes leads to decreased ovarian size via ceased folliculogenesis. **A)** Average size of ovary of CDK12^+/+^ and CDK12^-/-^ females. The number of females for each genotype is shown in parentheses. Data are presented as mean ± SE; Student’s t-test: ***p < 0.001. **B)** The ovary of the CDK12^-/-^ forms a corpus luteum (CL). Representative sections of histologic structures in ovaries of Cdk12-deficient females (CDK12^-/-^) and CDK12^+/+^; scale bar 400 µm.

**SI Fig. 3:**
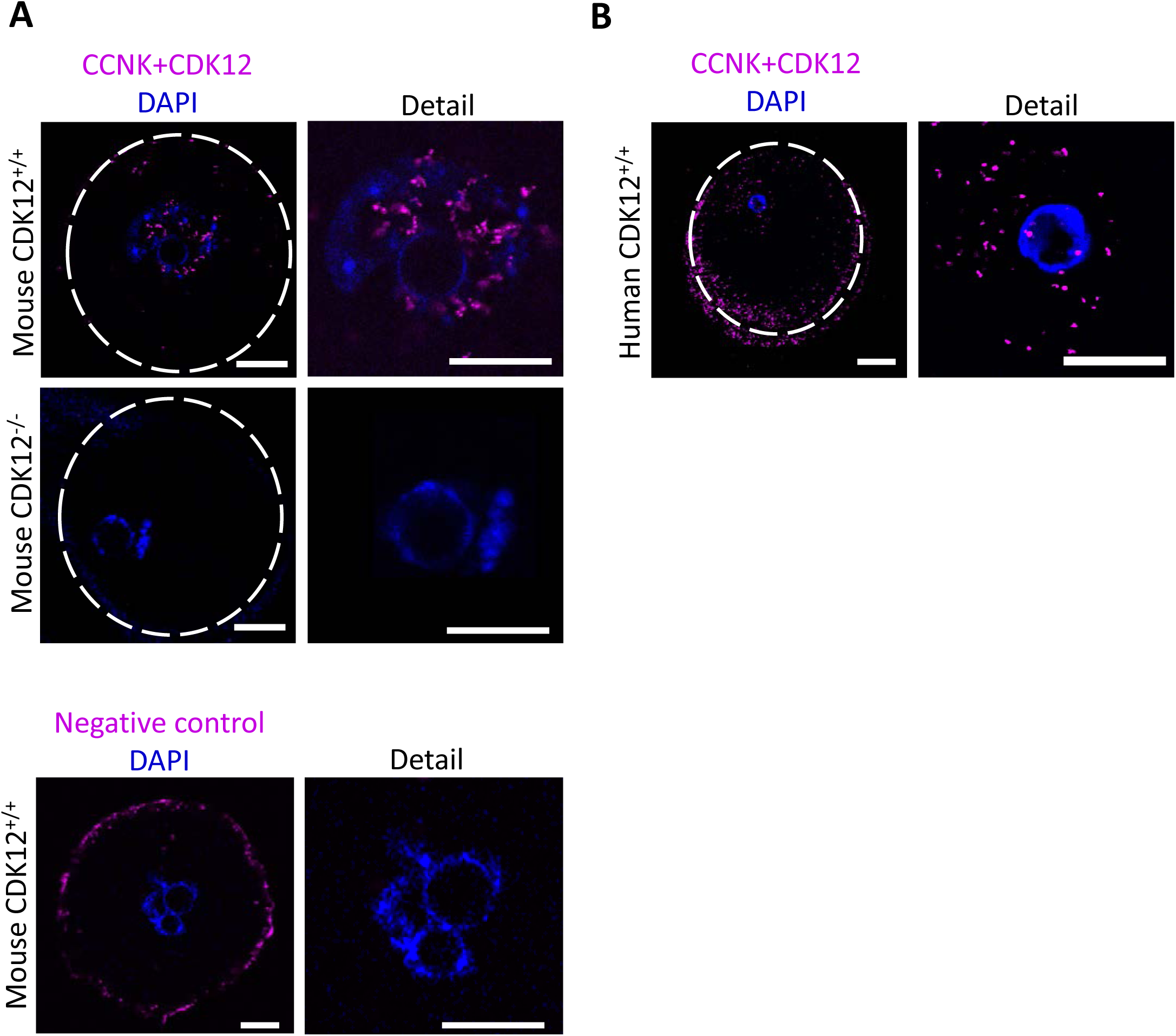
The CDK12/CCNK complex is localized to the nucleus of the oocyte in mice and humans. **A)** Representative images of proximity ligation assay showing the nuclear localization and physical interaction of CDK12 and CCNK (purple) in mouse CDK12^+/+^ GV oocytes and no interaction in CDK12^-/-^ oocytes. Detail shows higher magnification of the nuclear region; DAPI (blue); the dashed line shows the cell cortex. Representative images are from three independent biological replicates; n ≤28; scale bars 20 µm. **B)** Representative images of proximity ligation assay showing the nuclear localization and physical interaction of CDK12 and CCNK (purple) in human oocytes. Detail shows higher magnification of the nuclear region; DAPI (blue); the dashed line shows the cell cortex. Representative images are from three independent biological replicates; n=8; scale bar 20 µm.

**SI Fig. 4:**
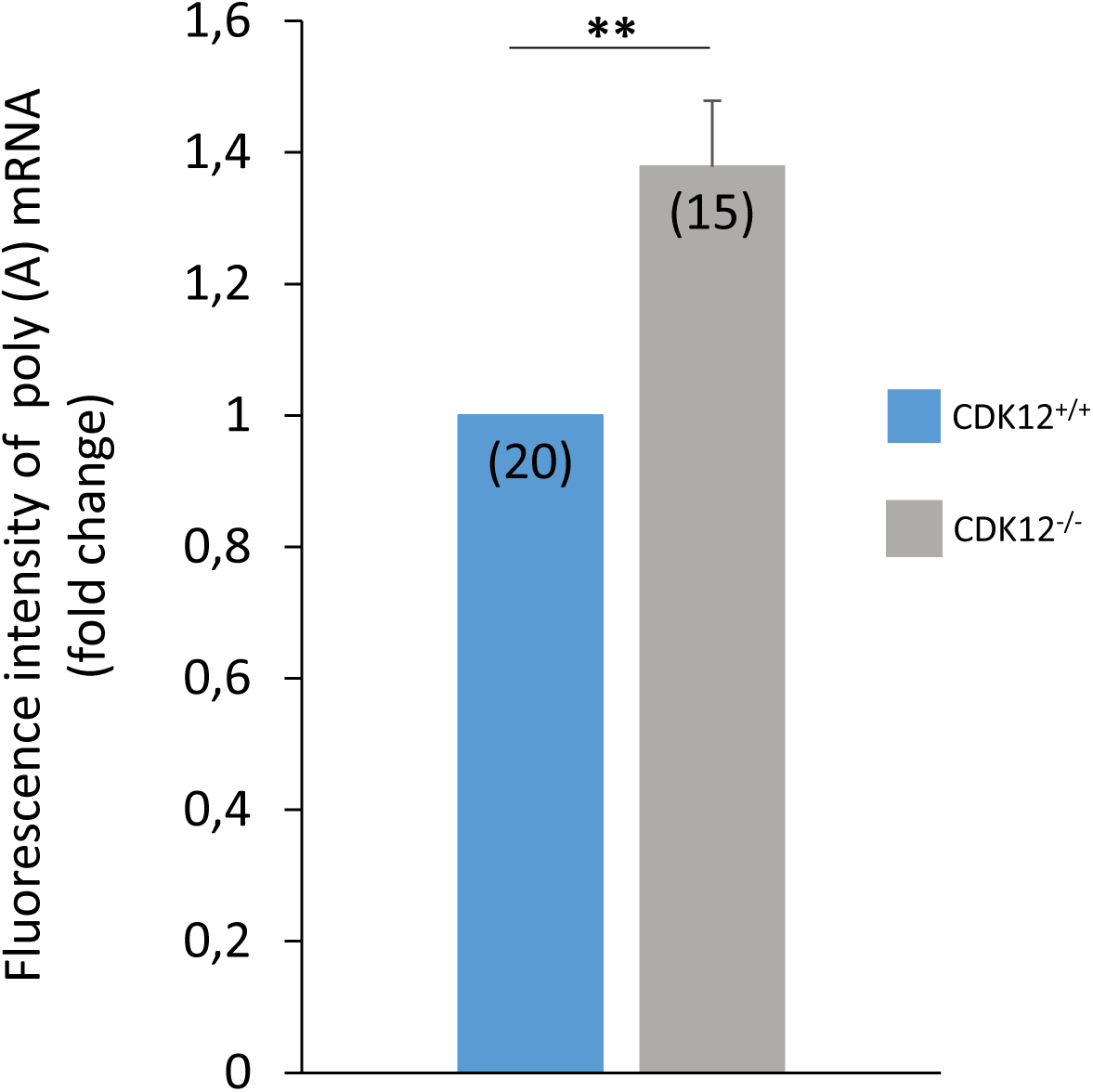
Global poly(A) RNA does not differ between oocytes of different genotypes. Quantification of poly(A) RNA fluorescence intensity in CDK12^+/+^ and CDK12^-/-^ oocytes from (Fig. 6D). The values from CDK12^+/+^ were set as 1. Data from three independent biological replicates, number of cells shown in parentheses. Data are presented as mean ± SE; Student’s *t*-test: **p < 0.01.

**SI Fig. 5:**
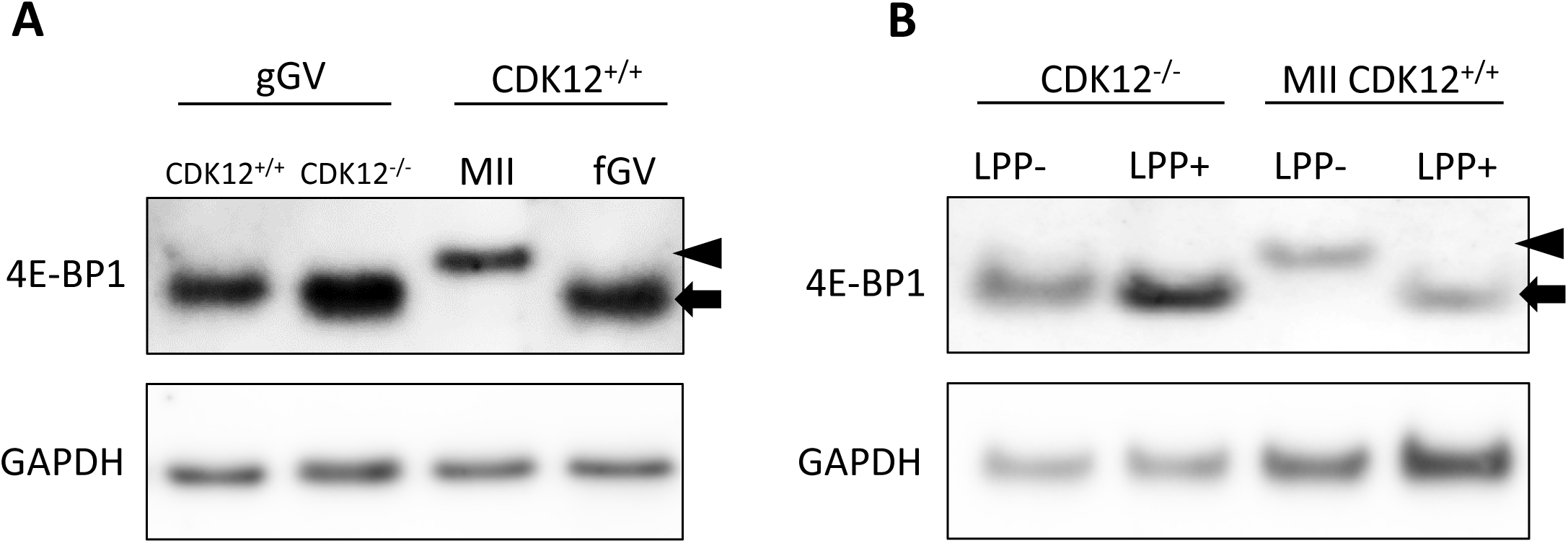
4E-BP1 is non-phosphorylated in growing oocytes. **A)** Western blot analysis of 4E-BP1 expression and phosphorylation shift in growing (gGV) oocytes from different genetic maternal sources. Non-phosphorylated (arrow) and phosphorylated (arrowhead) 4E-BP1. Mature (MII) and fully grown (fGV) oocytes from CDK12^+/+^ were used as controls. GAPDH was used as a loading control. Data were obtained from three biological replicates. **B)** Western blot analysis of phosphatase treatment (LPP+) of oocyte samples. Non-phosphorylated (arrow) and phosphorylated (arrowhead) form of 4E-BP1 in growing CDK12^-/-^ oocytes (gGV). Mature (MII) CDK12^+/+^ oocytes were used as a control. GAPDH was used as a loading control. Data are from two biological replicates.

**Supplementary Figure.**
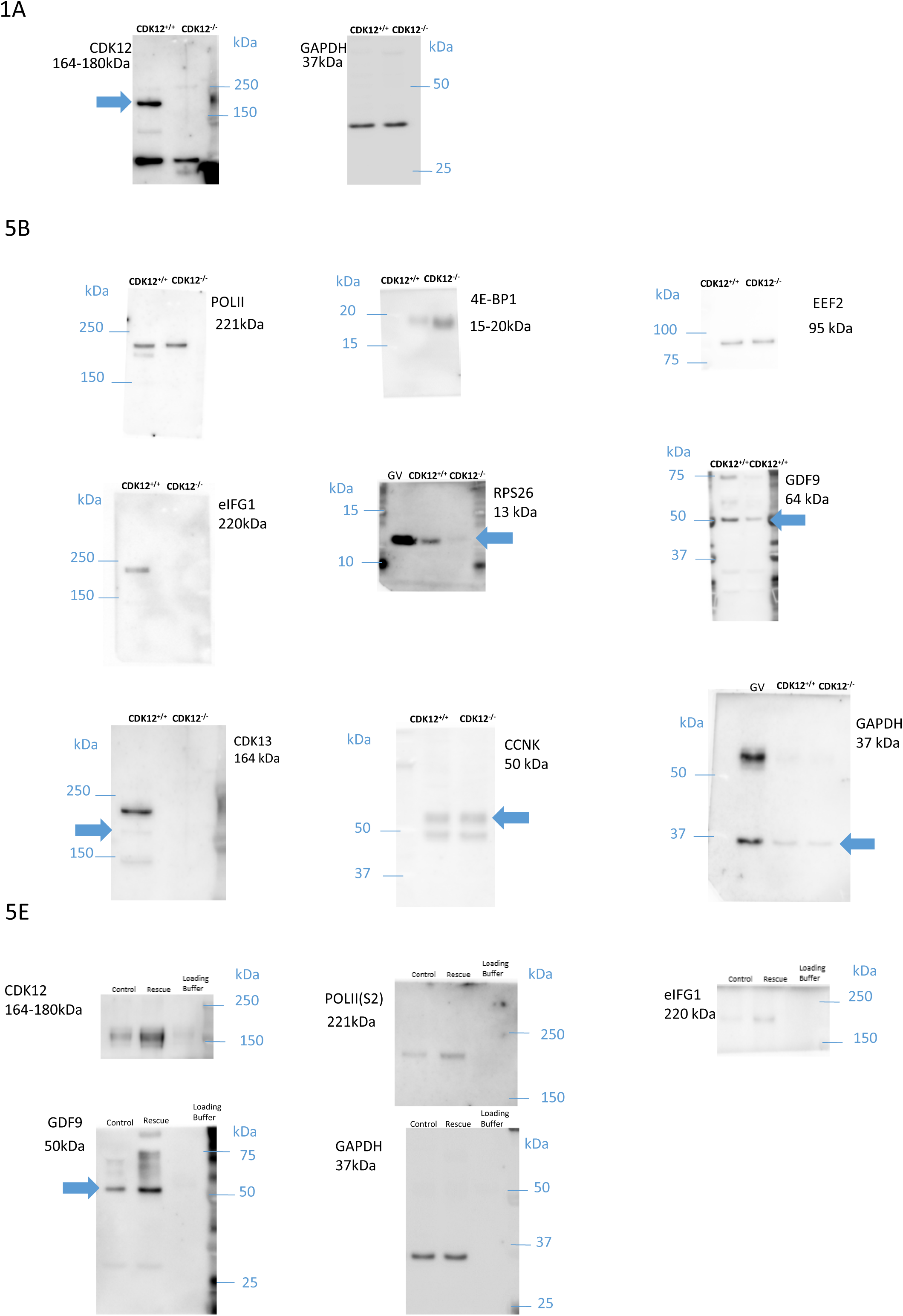
Full immunoblots of segments shown in the main Figure 1A and Figure 5B, 5E. Arrows denote the bands used.

**Supplementary Figure 1.**
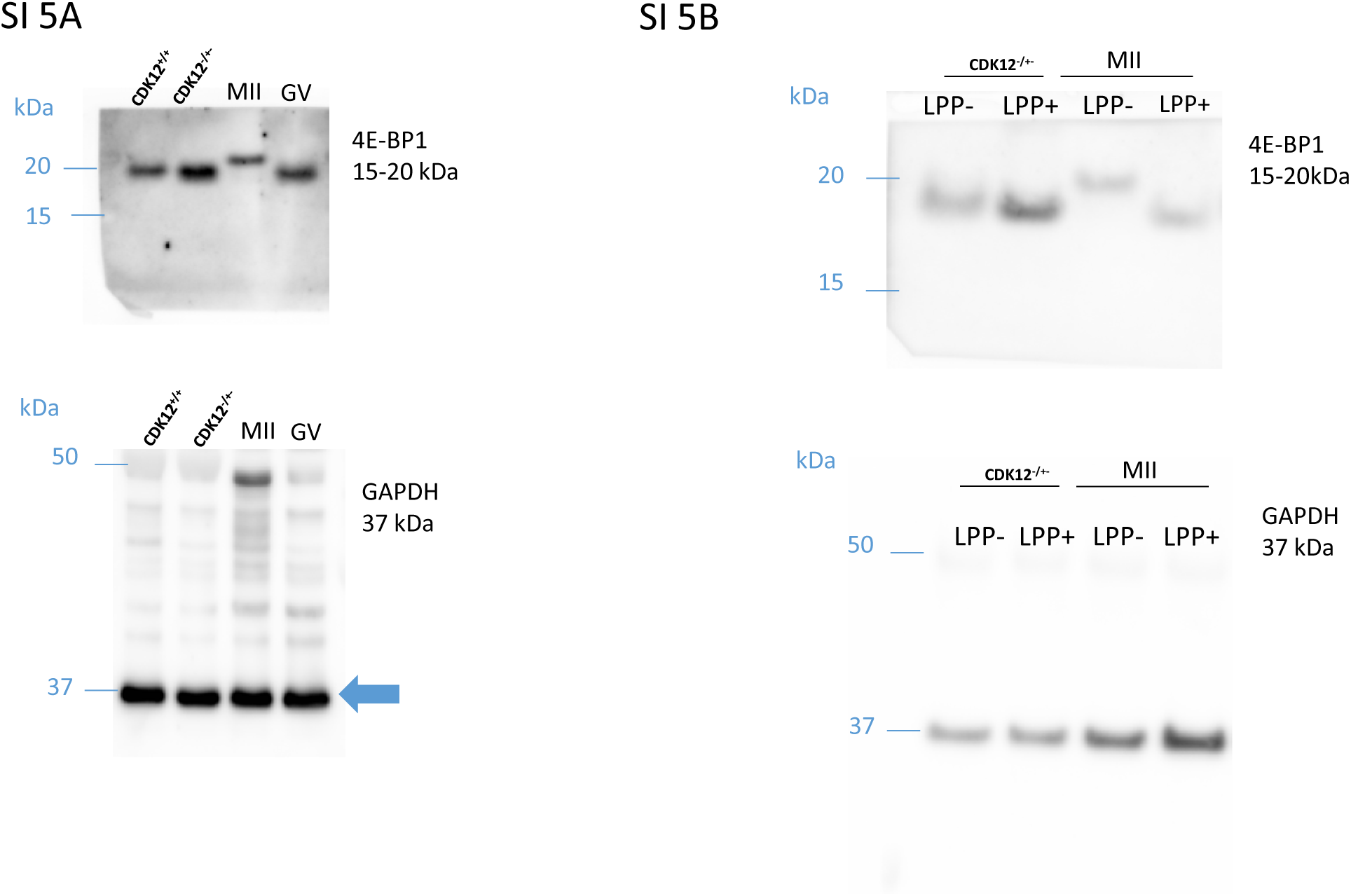
Full immunoblots of segments shown in SI

## Notes

### Competing Interest Statement

The authors have declared no competing interest.

